# A comparison of immunoglobulin IGHV, IGHD and IGHJ genes in wild-derived and classical inbred mouse strains

**DOI:** 10.1101/631754

**Authors:** Corey T. Watson, Justin T. Kos, William S. Gibson, Leah Newman, Gintaras Deikus, Christian E. Busse, Melissa Laird Smith, Katherine J.L. Jackson, Andrew M. Collins

## Abstract

The genomes of classical inbred mouse strains include genes derived from all three major subspecies of the house mouse, *Mus musculus*. We recently posited that genetic diversity in the immunoglobulin heavy chain (IGH) gene loci of C57BL/6 and BALB/c mice reflect differences in subspecies origin. To investigate this hypothesis, we conducted high-throughput sequencing of IGH gene rearrangements to document IGH variable (IGHV), joining (IGHJ), and diversity (IGHD) genes in four inbred wild-derived mouse strains (CAST/EiJ, LEWES/EiJ, MSM/MsJ, and PWD/PhJ), and a single disease model strain (NOD/ShiLtJ), collectively representing genetic backgrounds of several major mouse subspecies. A total of 341 germline IGHV sequences were inferred in the wild-derived strains, including 247 not curated in the International Immunogenetics Information System. In contrast, 83/84 inferred NOD IGHV genes had previously been observed in C57BL/6 mice. Variability among the strains examined was observed for only a single IGHJ gene, involving a description of a novel allele. In contrast, unexpected variation was found in the IGHD gene loci, with four previously unreported IGHD gene sequences being documented. Very few IGHV sequences of C57BL/6 and BALB/c mice were shared with strains representing major subspecies, suggesting that their IGH loci may be complex mosaics of genes of disparate origins. This suggests a similar level of diversity is likely present in the IGH loci of other classical inbred strains. This must now be documented if we are to properly understand inter-strain variation in models of antibody-mediated disease.

## INTRODUCTION

Inbred mouse strains are critical to biomedical research, and many of the most important strains such as DBA, C57BL, C3H, CBA and BALB/c have now been in use for almost a century ^1^. The C57BL and BALB/c strains have been particularly important for our understanding of the biochemistry and immunogenetics of immunoglobulins (IG) ^2–4^. This understanding was achieved despite a lack of detailed knowledge of the antibody genes of these and other inbred laboratory mouse strains, and until recently it was thought that the genes present in these different strains were likely to be highly similar ^5^.

Mouse antibody genes were first identified using cell lines derived from BALB/c mice because of the availability of mineral-oil induced plasmacytomas from this strain ^4^. The cataloguing of BALB/c antibody genes effectively ceased with the emergence of the C57BL/6 strain as the workhorse of transgenic and genomic studies. The IG heavy chain region (IGH) locus of the C57BL/6 strain was therefore the first to be sequenced and annotated ^6, 7^. Part of the IGH locus of the 129S1 strain was also reported ^8^, before comprehensive genomic investigation of mouse germline IGHV genes essentially ceased. By this time, two databases had catalogued mouse IGH genes and apparent allelic variants of these genes: the VBASE2 ^9^ and IMGT ^10^ databases. A positional nomenclature was then developed by IMGT, based upon the mouse genome reference sequence, while an alternative positional nomenclature was developed by Johnston and colleagues, based upon an alternative assembly of the C57BL/6 genome ^6^. A non-positional gene sequence identifier system was also developed by VBASE2 ^9^.

The study of IGH gene variation in humans followed a similar trajectory to that of the mouse. Genes and likely allelic variants were reported over a twenty-year period, starting in the late 1970s ^11^. There was a sharp decline in the reporting of new sequences once the complete human IGH locus was published in 1998 ^12^, but the advent of high-throughput sequencing of human antibody genes reawakened interest in the documentation of allelic variants ^13, 14^. A surprising level of antibody gene variation, including structural variation of the IGH locus, has since been shown within the human population ^15–18^, and such variation can have important consequences for the development of a suitable protective antibody repertoire ^19, 20^. A similar exploration of the immunoglobulin gene variation and antibody repertoires is now beginning in the mouse.

Many recently discovered allelic variants of IGH variable (IGHV), diversity (IGHD), and joining (IGHJ) genes have been identified from sets of VDJ gene rearrangements, using a process of inference ^21^. Rearranged VDJ sequences are often affected by somatic hypermutations, and such mutations are distributed throughout VDJ rearrangements. When the same mismatch to a known germline gene is repeatedly observed in data from a single subject, however, it is more likely that the nucleotide in question is a single nucleotide variant (SNV), rather than being a nucleotide that has arisen by somatic hypermutation ^21^. When such mismatches are repeatedly seen in a large set of VDJ rearrangements having diverse CDR3 regions and amplified from a single individual, the inference of a previously undiscovered gene polymorphism may be made with confidence. The discovery of allelic variants by inference is now a feature of many human repertoire studies, and this is facilitated by a number of recently developed utilities ^22–25^. When this approach was applied to the mouse, the outcome was quite unexpected.

Analysis of thousands of C57BL/6 VDJ rearrangements identified 99 of the 114 germline IGHV genes that have been reported to be functional in this strain ^5^. It was concluded that the remaining 16 genes are either non-functional or are expressed at such low frequencies that they would only be detectable in deeper sequencing studies. It was also concluded that all IGHV genes carried by any strain of inbred mouse that are expressed at moderate frequencies should be readily determinable by inference from VDJ gene datasets.

An analysis of BALB/c VDJ rearrangements was then performed, and 163 BALB/c IGHV gene sequences were identified ^5^, only half of which were present in the IMGT database ^10^. The expression of ten unique IGHD sequences and four IGHJ genes was confirmed, while three other reported BALB/c IGHD genes appeared to be non-functional. Although the identification of BALB/c IGHV genes was almost certainly incomplete, the study successfully captured the germline gene variability that accounts for almost all of the genetic variation in the repertoire of rearranged heavy chain genes.

These were not the first striking differences seen in the IGH loci of BALB/c and C57BL/6 mice. Historically, these strains have been known to carry different IGH constant region gene haplotypes, Igh-1^a^ and Igh-1^b^ respectively, and in the past these haplotypes were determined serologically. Sequencing studies later showed striking differences between the two haplotypes, particular at the IGHG2 gene locus. Controversy has surrounded the evolutionary origins of this allotypic variation, but genomic evidence led to the suggestion that it resulted from gene duplication and gene loss in different mouse subspecies ^26^.

The differences, seen by Collins and colleagues ^5^, between the germline BALB/c and C57BL/6 IGHV genes were so profound that it was proposed that the IGH genes of these strains may have had their origins in different subspecies of the house mouse. Three major subspecies of the house mouse have been described (*Mus musculus musculus*, *M. m. domesticus* and *M. m. castaneus*) and subsequent analysis of genomic SNV data supported the origins of BALB/c and C57BL/6 mice from *M. m. domesticus* and *M. m. musculus* respectively ^27^ (see Table 1 and Supplementary figure 1).

**Table 1:** The intended *Mus musculus* subspecies origins of inbred strains of mice, and the subspecies identity of the IGH loci of inbred strains, as determined by SNV analysis.

| Strain | Subspecies Origin | Subspecies identity by SNV analysis |
| --- | --- | --- |
| PWD/PhJ | *M. m. musculus* | *M. m. musculus* |
| LEWES/EiJ | *M. m. domesticus* | *M. m. domesticus* |
| CAST/EiJ | *M. m. castaneus* | *M. m. castaneus* |
| MSM/MsJ | *M. m. mollosinus* | *M. m. musculus* |
| C57BL/6 | N/A | *M. m. musculus* |
| BALB/c | N/A | *M. m. domesticus* |
| NOD/ShiLtJ | N/A | *M. m. musculus* |

To test the hypothesis that BALB/c and C57BL/6 IGH genes are derived from different subspecies of the house mouse, in this study we first document IGHV, IGHD and IGHJ germline genes in a number of wild-derived inbred mouse strains representing each of the three major subspecies of the house mouse (CAST/EiJ: *M. m. castaneus*; PWD/PhJ: *M. m. musculus*; LEWES/EiJ: *M. m. domesticus*), and of a wild-derived strain (MSM/MsJ) that originated from *M. m. molossinus* mice that are generally considered to be hybrids of *M. m. musculus* and *M. m. castaneus* (see Table I). We then compare these genes to those of the C57BL/6 and BALB/c strains, in order to infer the ancestry of these classical inbred strains. The wild-derived strains used in this study were developed in the 1970s from pairs of wild mice from known locations, with the intention that each inbred strain would carry a genome that was derived from a single subspecies of the house mouse. To explore the differences between strains that appear, based upon preliminary SNV analysis, to be derived from the same subspecies of the house mouse, we also investigated the NOD/ShiLtJ inbred strain (Table 1 and Supplementary figure 1).

Although it was not possible in this study to document all rearrangeable genes of the IGH loci, coverage was sufficient to allow broad conclusions to be drawn. Unexpectedly, the NOD and C57BL/6 gene IGHV, IGHD, and IGHJ loci appear to be almost exactly the same, while striking differences exist between each wild-derived strain and both the C57BL/6 and BALB/c strains. The divergence that was seen between the strains suggests that the IGH loci of inbred mouse strains are likely to harbor so much genetic variation that if the antibody responses of these and other classical inbred mouse strains are to be properly understood, it will be essential to fully document their germline genes. The divergence between the strains also suggests that a single positional mouse gene nomenclature based upon the C57BL/6 genome reference sequence may fail to properly represent the genes of many important inbred mouse strains.

## RESULTS

### Defining inferred IGH germline gene sets in diverse inbred mouse strains

To ensure the highest quality input data, prior to VDJ assignment we leveraged the PacBio circular consensus (CCS2) algorithm to generate high quality circular consensus reads for each sample. The average read length across the libraries sequenced was 22.5 Kb (Supplementary table 1). The long read lengths paired with the target amplicon library size of ∼1200 bp, resulted in a mean of 23.9 circular consensus passes per amplicon (Supplementary table 1). Finally, we applied a Q30 cutoff to data from each library, resulting in a total of 36782, 43522, 43173, 28136 and 43044 pre-mapped raw reads for CAST/EiJ, LEWES/EiJ, MSM/MsJ, PWD/PhJ, and NOD/ShiLtJ, respectively, with mean CCS read scores of 1 (Supplementary table 1). Each of these high quality read datasets were used for IGHV, IGHD, and IGHJ gene assignment, clonal assignment, and germline gene inference. Read data for each strain have been submitted to the Sequence Read Archive (SRA) under the BioProject ID PRJNA533312.

The discovery stage of our inference pipeline for each strain was modelled after that used previously ^5^ (see Materials and Methods below). Clonal assignment and clustering resulted in a total of 5261 (CAST/EiJ), 5042 (LEWES/EiJ), 4374 (MSM/MsJ), 3827 (PWD/PhJ), and 3743 (NOD/ShiLtJ) unique clones; a single representative sequence from each of the identified clones was then randomly selected to use for germline inference. With these sequences as input, a total of 87, 78, 84, 92, and 84 germline IGHV sequences were inferred for CAST/EiJ, LEWES/EiJ, and MSM/MsJ, PWD/PhJ, and NOD/ShiLtJ, respectively (Figure 1a; Supplementary table 2). Each inferred germline sequence was represented by at least 0.1% of the total clones observed in a given strain; the numbers of clones representing each inference are provided in Supplementary table 2 and plotted in Supplementary figure 2.

**Figure 1.**
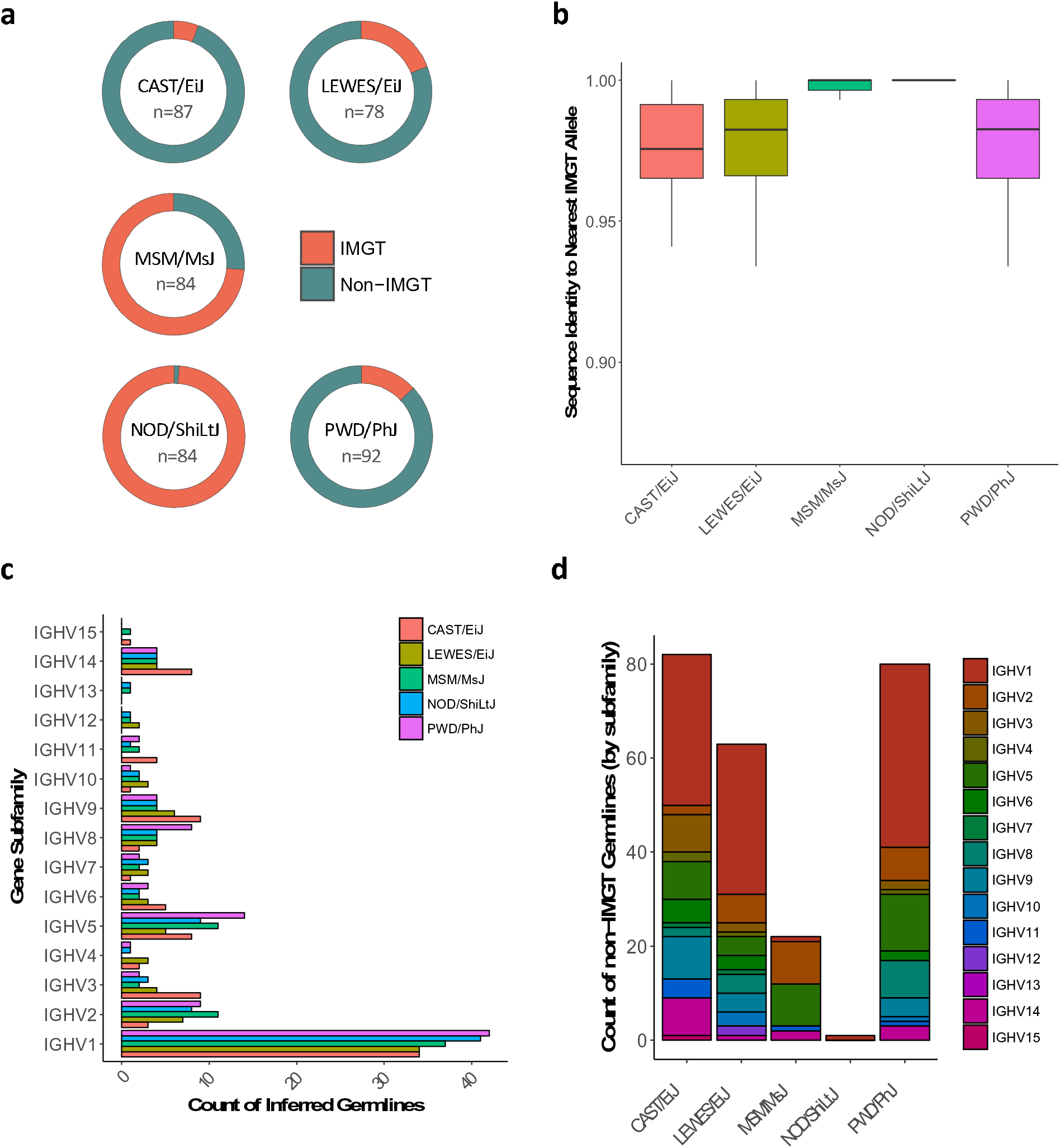
Comparisons of inferred germline IGHV sequences from each strain (CAST/EiJ, n=1; LEWES/EiJ, n=1; MSM/MsJ, n=1; PWD/PhJ, n=1; NOD/ShiLtJ, n=1) to those represented in the mouse IMGT database (imgt.org). (**a**) Donut plots depicting the proportion of inferred germline sequences from each strain that align to a known IMGT allele with 100% match identity. (**b**) Boxplots depicting the sequence similarities of inferred germline sequences from each strain when compared to the closest known IMGT allele. (**c**) The count of identified germline sequences from each strain representing known mouse IGHV gene subfamilies. (**d**) The count of non-IMGT inferred germline sequences in each strain, partitioned by IGHV gene subfamily.

Across the wild-derived strains, the validity of some inferred germline IGHV sequences was supported by their presence in the IMGT mouse reference directory (http://www.imgt.org) (Figure 1a). This included 5 sequences in CAST/EiJ, 15 in LEWES/EiJ, 62 in MSM/MsJ, and 12 in the PWD/PhJ strain. Sequences inferred from the wild-derived strains were, however, dominated by non-IMGT alleles (247/341, 72%; Figure 1a). Of special note, all but one of the NOD/ShiLtJ gene sequences have previously been reported in the IMGT reference directory, and all of these 83 sequences are reported there as C57BL/6 sequences.

Additional supporting evidence was found in other public sequence repositories for 44 non-IMGT sequences (Supplementary table 2). Two non-IMGT sequences were found in the MSM/MsJ strain with perfect matches to sequences reported in either VBASE2 (http://www.vbase2.org/) or the NCBI reference set. This was also true for 2, 29, and 11 non-IMGT sequences inferred from CAST/EiJ, LEWES/EiJ, and PWD/PhJ, respectively.

The sets of germline inferences were used as starting databases for analysis by IgDiscover ^22^, to seek further validate the inferences. This analysis confirmed 93% of the inferences: 82/87 (94%), 68/78 (87%), 76/84 (90%), 86/92 (93%), and 84/84 (100%) of the inferred sequences for CAST/EiJ, LEWES/EiJ, and MSM/MsJ, PWD/PhJ, and NOD/ShiLtJ, respectively.

Most of the inferences in the wild-derived strains that were not confirmed by IgDiscover were present at low copy number (Supplementary table 2 and Supplementary figure 3), though a few unconfirmed sequences were seen at relatively high copy number, including CAST-IGHV9-2 (28 clones), LEWES-IGHV3-2 (25 clones) and MSM-IGHV2-4 (50 clones). Many of the unconfirmed sequences were supported by other secondary evidence from public databases, or were also seen in other strains from this study (Supplementary table 2). This included seven sequences that are present among whole-genome shotgun sequence data generated by the Mouse Genome Project (https://www.sanger.ac.uk/science/data/mouse-genomes-project). Just 11 of the sequences unconfirmed by IgDiscover lacked additional evidence supporting their validity, representing only ∼2.5% of the total number of sequences identified in our dataset. Issues of false negatives by IgDiscover, and other related inference tools, have been reported, but have not been fully explained ^24^. As a consequence, after additional manual review of our results, the full sets of inferences were retained and are reported here.

In some cases, novel sequences from wild-derived strains were quite divergent from published IGHV sequences, varying from 87.02% to 99.66% sequence identity. This was also dependent on strain in that, among the wild-derived strains, inferred germlines from CAST/EiJ, LEWES/EiJ, and PWD/PhJ exhibited much greater sequence divergence from IMGT alleles than sequences inferred in MSM/MsJ (Figure 1b).

The counts of genes within the different IGHV families were generally comparable across the five strains (Figure 1c), with some exceptions. The germline repertoires of all strains were clearly dominated by the IGHV1 family. However, the numbers of IGHV genes in other subgroups were more variable. For example, the repertoire of CAST/EiJ harbored fewer IGHV2 genes relative to the other strains, but greater numbers of IGHV3, IGHV6, IGHV9, and IGHV14 genes. At least one representative germline sequence of subfamilies IGHV1-IGHV3, IGHV5-IGHV10, and IGHV14 was inferred from all strains; in contrast, sequences of the remaining subfamilies, which are all small subfamilies in the C57BL/6 strain, were absent in at least one strain. IGHV13 and IGHV15 sequences were only observed in two of the five strains. Whether this represents a genuine lack of functional IGHV subfamily sequences in these strains (e.g. as a result of pseudogenization or genomic deletion), or whether this is due to under sampling of these repertoires is not clear. Deeper sequencing and ultimately comprehensive genomic characterization in each strain would be needed to fully assess this. Consistent with the general subfamily distributions observed, the majority of non-IMGT sequences in CAST/EiJ (n=32), LEWES/EiJ (n=32), and PWD/PhJ (n=39) were represented by IGHV1 genes (Figure 1d). In contrast, however, although the MSM/MsJ repertoire was also dominated by IGHV1, the majority of non-IMGT sequences identified in that strain were from IGHV2 (n=9) and IGHV5 (n=9) (Figure 1d).

Analysis was also performed to identify previously unreported polymorphism of IGHD and IGHJ genes, and to define the sets of IGHD and IGHJ genes in each mouse strain. Analysis of IGHD gene alignments led to the identification of eight or nine IGHD genes in each of the strains, with four novel sequences being identified as likely allelic variants of IGHD1-1 (n=2), IGHD2-3 (n=1) and IGHD2-12 (n=1) (Figure 2a and Supplementary table 3). These polymorphisms could be determined with absolute certainty because the critical nucleotides that distinguish them from previously reported IGHD sequences are sufficiently distant from the IGHD gene ends. No variants were identified with differences in the 5’ or 3’ terminal nucleotides of the IGHD genes, but we cannot exclude the possibility that our analysis failed to detect such variation.

**Figure 2.**
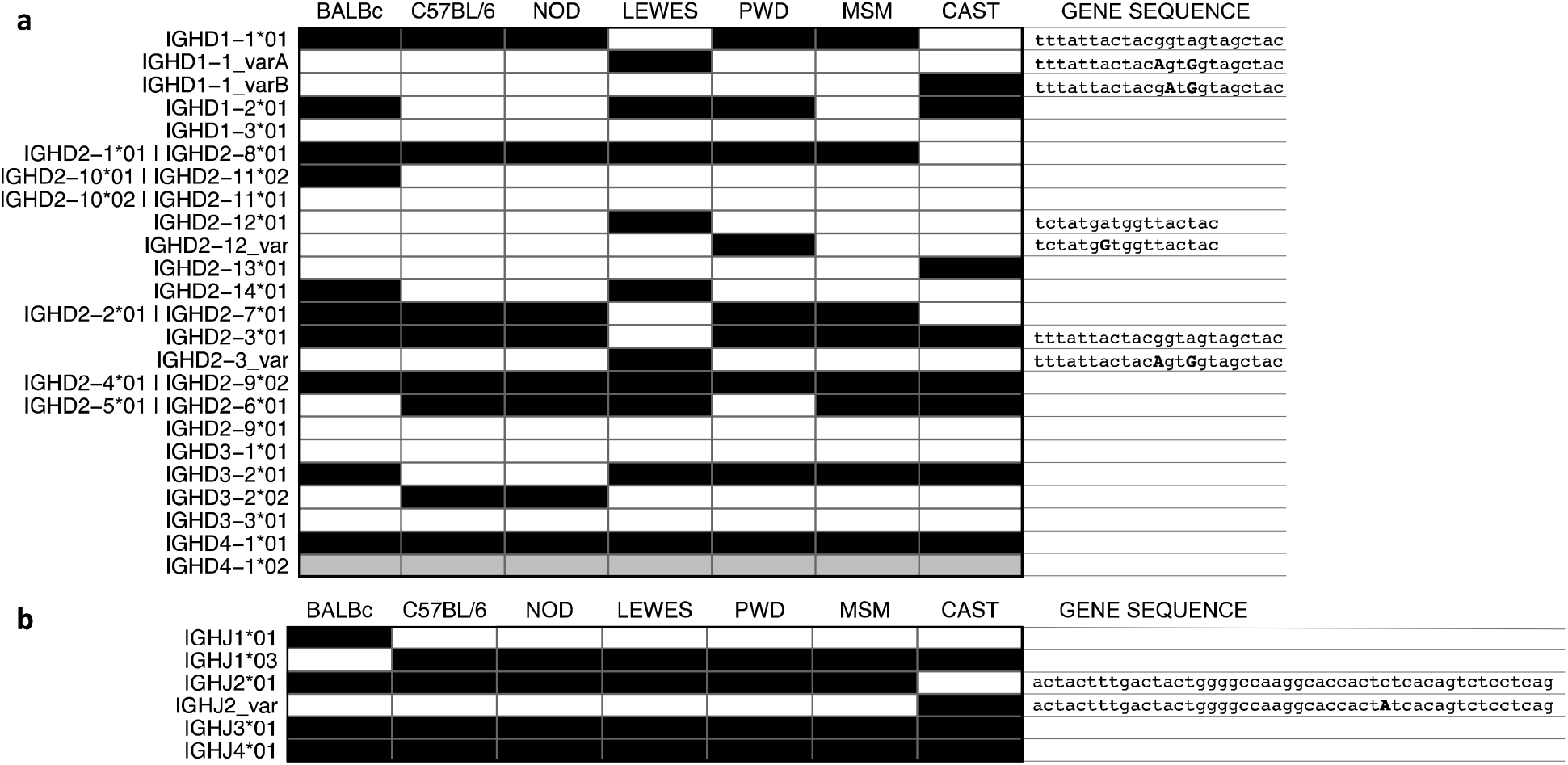
IGHD and IGHJ inferred germline sequences among mouse strains (CAST/EiJ, n=1; LEWES/EiJ, n=1; MSM/MsJ, n=1; PWD/PhJ, n=1; NOD/ShiLtJ, n=1). (**a**) Tile plot depicting the presence of IGHD genes across the different mouse strain. Filled black cells indicate the presence of the gene. For novel variants, the sequences are listed along with the *01 reference allele, with nucleotide differences for the variant indicated in uppercase bold. (**b**) Tile plot depicting the presence of IGHJ genes across the different mouse strains. Black cells indicate that a gene was confirmed as present in a strain, white indicates that a gene wasn’t confirmed for a strain and gray indicates genes whose presence remains uncertain. For the novel variant, the sequence is listed along with the *01 reference allele, with nucleotide differences for the variant indicated in uppercase bold.

The IGHD genes of NOD/ShiLtJ were shown to be identical to those of the C57BL/6 mouse. The MSM/MsJ and PWD/PhJ strains share 6 IGHD genes with the C57BL/6 strain, and carry two and three additional unique genes respectively. The PWD/PhJ strain expresses a variant of the 26 nucleotide IGHD2-12 gene (Figure 2a and Supplementary table 3). The LEWES/EiJ strain shares five sequences with the BALB/c strain, but also expresses four additional sequences, including a previously unreported variant of the IGHD1-1 gene. This variant differs from the IGHD1-1*01 sequence at two positions (Figure 2a and Supplementary table 3). The CAST/EiJ strain also expresses a novel IGHD1-1 variant, distinct from that observed in LEWES/EiJ, as well as six genes that are expressed by other strains in the study. Only CAST/EiJ expresses the IGHD2-13*01 gene. The IGHD4-1*01 and IGHD4-1*02 are highly similar gene segments sharing 9 of 11 and 9 of 10 nucleotides, respectively. In all strains, full length examples of both IGHD4-1 variants are observed, but the shorter IGHD4-1*02 is difficult to confirm with confidence as such a sequence can be derived by trimming 2 nucleotides from the 5’ end of *01, followed by the non-template addition of ‘c’.

All strains were found to carry four functional IGHJ genes, which were each used at sufficient frequency to make their identification unequivocal (Figure 2b and Supplementary table 4). Only in CAST/EiJ was there evidence of a novel allele (IGHJ2_var). This novel allele harbored a single nucleotide difference from the closest known allele in the IMGT reference directory (Figure 2b). The other CAST/EiJ IGHJ genes were shared with the other strains and were identical to the genes carried by C57BL/6 mice. No strain carried the BALB/c IGHJ1*01 allele ^5^.

### Extensive IGHV germline diversity and limited overlap between strains

We next investigated the extent of overlap of IGHV sequences among the surveyed strains. The sets of germline genes inferred from each inbred strain were compared to sequences identified in all other strains to determine how many sequences were identical between strains. Comparisons were additionally made with previously published inferences from BALB/c mice (n=163) ^5^, and with the IMGT repertoire of functional C57BL/6 sequences (n=114). Surprisingly little overlap was observed between strains, and the majority of inferred germline sequences were unique to a single strain (Figure 3a). Among the wild-derived strains surveyed, CAST/EiJ had the highest number of unique germline sequences (76/87 sequences). On the other hand, 83 shared sequences were observed in NOD/ShiLtJ and C57BL/6, with 48 of these 83 sequences being additionally shared by MSM/MsJ. A single sequence was identified in five different strains (LEWES/EiJ, NOD/ShiLtJ, CAST/EiJ, PWD/PhJ, and C57BL/6; Supplementary table 2).

**Figure 3.**
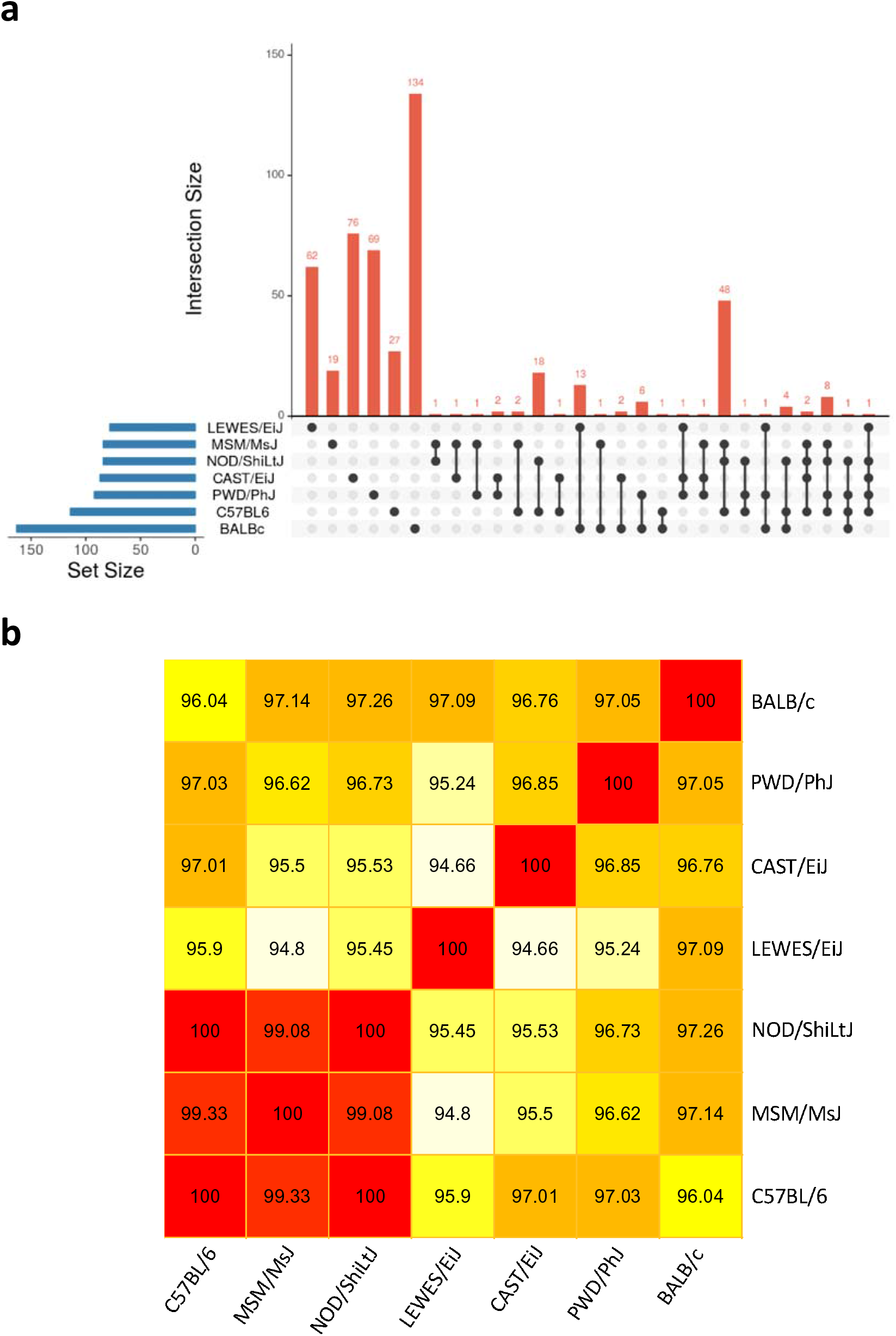
Relationships of IGHV inferred germline sequences among mouse strains (CAST/EiJ, n=1; LEWES/EiJ, n=1; MSM/MsJ, n=1; PWD/PhJ, n=1; NOD/ShiLtJ, n=1). **(a)** Upset plot depicting the size of the germline set from each of the analyzed strains (left), as well as the numbers of sequences either unique to a given strain or shared among strains (identical sequences). (**b**) Heatmap depicting the mean percent sequence match identities among inferred IGHV germline sets for each pair-wise strain comparison (see also Supplementary figure 4).

We further explored interstrain IGHV sequence relationships by estimating the average sequence similarities of IGHV sets between strains. Consistent with sequence overlaps presented in Figure 3a, we noted a range of mean pairwise sequence identities, depending on the strains in question. For example, sequences in MSM/MsJ, C57BL/6, and NOD/ShiLtJ, strains which share the most identical sequences with one another (Figure 3a), also have high average pairwise sequence identities (>99%; Figure 3b; see also Supplementary figure 4 for full pairwise comparisons). This is in contrast to mean identities observed for all other pairwise strain comparisons, which ranged from 94.8% to 97.1%. These levels of identity also generally matched the number of shared sequences between strains. For example, LEWES/EiJ shared the most identical sequences with BALB/c, and among all pairwise sequence comparisons between LEWES/EiJ and other strains, the highest mean sequence identity was with the BALB/c germline set (97.1%; Figure 3b).

No attempt was made in this study to determine whether or not any pairs of highly similar sequences could be allelic variants of a single gene. In light of known structural variation in the IGH loci of C57BL/6 and BALB/c mice ^7^, we must assume that significant structural variation is possible between the loci of the wild-derived strains reported here. In a locus where gene duplications and deletions, as well as pseudogenization, are central drivers of evolution, and where many sets of highly similar genes are found in each strain, the documentation of allelic variants of any gene will require comprehensive genomic sequencing.

### Implications of inferred germline references on repertoire alignments

The re-analysis of the VDJ datasets from each strain using the strain-specific inferred repertoires dramatically impacted upon the repertoire-level alignment metrics (Table 2). This was especially true for CAST/EiJ which was most poorly represented in the IMGT reference directory. Considering the IgBLAST output for the two different repertoires, just 3.8% of IGHVs and 49.4% of IGHJs were unmutated if the IMGT reference directory was used for analysis. This increased to 87.8% and 88.2% for IGHV and IGHJ respectively using the CAST/EiJ inferred germline references. LEWES/EiJ and PWD/PhJ also experienced significant improvements for the IGHV, shifting the frequency of unmutated reads within the IgM repertoires from 35.6 to 88.6% and 13.9 to 82.9%, respectively.

**Table 2:** IgBLAST results using the IMGT database compared to novel strain-specific repertoires characterized in the current study.

|  |  |  | IGHV |  |  |  | IGHD |  |  |  | IGHJ |  |  |  |
| --- | --- | --- | --- | --- | --- | --- | --- | --- | --- | --- | --- | --- | --- | --- |
|  |  |  | IGHV (n) * | unmutated IGHVs (n) ^ | unmutated reads (%) # | mean SHM (%) | IGHD (n) * | unmutated IGHDs (n) ^ | unmutated reads (%) # | mean SHM (%) | IGHJ (n) * | unmutated IGHJs (n) ^ | unmutated reads (%) # | mean SHM (%) |
| IgBLAST | IMGT | CAST | 109 | 30 | 3.8% | 2.21% | 16 | 16 | 87.9% | 0.92% | 8 | 4 | 49.4% | 1.45% |
|  |  | LEWES | 97 | 27 | 35.6% | 1.59% | 16 | 16 | 90.4% | 1.17% | 7 | 7 | 88.8% | 0.52% |
|  |  | MSM | 110 | 86 | 71.3% | 0.52% | 17 | 17 | 98.3% | 0.12% | 7 | 6 | 90.7% | 0.39% |
|  |  | NOD | 98 | 96 | 82.7% | 0.10% | 16 | 15 | 96.4% | 0.22% | 6 | 5 | 93.2% | 0.18% |
|  |  | PWD | 121 | 33 | 13.9% | 1.91% | 17 | 17 | 96.8% | 0.16% | 6 | 5 | 89.3% | 0.45% |
| IgBLAST | inferred | CAST | 87 | 87 | 87.8% | 0.25% | 9 | 9 | 96.2% | 0.30% | 4 | 4 | 88.2% | 0.49% |
|  |  | LEWES | 78 | 78 | 88.6% | 0.20% | 11 | 11 | 97.8% | 0.18% | 4 | 4 | 88.3% | 0.53% |
|  |  | MSM | 84 | 84 | 88.4% | 0.25% | 9 | 9 | 98.3% | 0.12% | 4 | 4 | 90.2% | 0.39% |
|  |  | NOD | 84 | 84 | 82.9% | 0.14% | 9 | 9 | 97.6% | 0.16% | 4 | 4 | 93.1% | 0.18% |
|  |  | PWD | 92 | 92 | 82.9% | 0.32% | 10 | 10 | 97.2% | 0.19% | 4 | 4 | 88.8% | 0.45% |
| * number of germline genes from reference |  |  |  |  |  |  |  |  |  |  |  |  |  |  |
| ^ number of germline genes from reference with 0 mismatches |  |  |  |  |  |  |  |  |  |  |  |  |  |  |
| # percent of reads assigned to unambiguous germline reference with 0 mismatches |  |  |  |  |  |  |  |  |  |  |  |  |  |  |

## DISCUSSION

This study was undertaken to investigate the hypothesis that IGH genes of subspecies of the house mouse are highly divergent, and to trace the subspecies origins of the IGH loci of BALB/c and C57BL/6 inbred mice. Since the mice that were used to establish the classical inbred strains of laboratory mice came from diverse and usually undocumented sources, we reasoned that differences in the loci of the various subspecies of the house mouse could explain the marked differences that were previously reported between the sets of IGHV genes found in the BALB/c and C57BL/6 strains. To test this hypothesis, we inferred the germline IGHV, IGHD and IGHJ genes of wild-derived strains representing each of the three major subspecies of the house mouse (CAST/EiJ: *M. m. castaneus*; PWD/PhJ: *M. m. musculus*; LEWES/EiJ: *M. m. domesticus*), and of a wild-derived strain (MSM/MsJ) that originated from *M. m. molossinus* mice that are generally considered to be natural hybrids of *M. m. musculus* and *M. m. castaneus* (see Table I). Whereas SNV-inferred haplotypes reported by Yang and colleagues ^28^ supported the reported subspecies origins of CAST/EiJ, PWD/PhJ, and LEWES/EiJ, these data suggested that the MSM/MsJ IGH locus is *M. m. musculus*-derived, with a SNV profile that is little different to that of the C57BL/6 mouse (see Table I and Supplementary figure 1). The IGH locus of the MSM/MsJ strain was therefore investigated in the hope that it would shed light on genetic variation within the loci of *M. m. musculus*-derived strains. For the same reason, the NOD/ShiLtJ strain was also included in this study, because SNV analysis suggested that it too carries a C57BL/6-like IGH haplotype.

In general, among the wild-derived strains surveyed in this study we observed surprisingly little overlap between IGHV sequences. While this was expected in comparisons of strains predicted to carry IGH loci originating from different subspecies, intriguingly there were notable differences in IGHV sequence sets between strains carrying loci of the same predicted subspecies. For example, PWD/PhJ mice only shared 13 (∼14%) IGHV gene sequences with the C57BL/6, NOD/ShiLtJ, and MSM/MsJ strains, despite the fact that all of these strains were predicted to share IGH loci of *M. m. musculus* origin. In fact, the PWD/PhJ strain shared almost the same number of IGHV sequences with the CAST/EiJ, LEWES/EiJ, and BALB/c strains (predicted to represent the *M. m. castaneus, M. m. domesticus* and *M. m. domesticus* subspecies respectively). PWD/PhJ IGHV sequences were also no more similar to sequences of other *M. m. musculus*-derived strains than to *M. m. castaneus*- or *M. m. domesticus*-derived strains. Similarly, although SNV analysis suggested that BALB/c and LEWES/EiJ mice both carry *M. m. domesticus*-derived IGH loci, only 13 (∼16%) of the identified LEWES/EiJ IGHV sequences are shared with the BALB/c strain. LEWES/EiJ IGHV sequences were collectively most similar to BALB/c genes, relative to the other strains sequenced here, but it is difficult to believe that the IGH loci of both strains are derived in their entirety from shared *M. m. domesticus* ancestors.

The IGHV gene sequences of the MSM/MsJ strain were particularly surprising. Few of the 84 MSM sequences were seen in any of the other three wild-derived strains. As expected, none of these sequences were amongst the 78 IGHV genes identified in the *M. m. domesticus*-derived LEWES/EiJ strain; however, there was also little identity with sequences identified in either of the subspecies that are said to have given rise to the hybrid *M. m. molossinus* mice. Only 10 of the 84 MSM/MsJ IGHV sequences matched those seen in the *M. m. musculus*-derived PWD/PhJ strain, and only 4 sequences matched those in the *M. m. castaneus*-derived CAST/EiJ strain. Instead, substantial identity was seen between MSM/MsJ mice and inbred C57BL/6 and NOD/ShiLtJ mice, that SNV analysis suggests are both *M. m. musculus*-derived (Supplementary figure 1). Genomic sequencing will be required to determine whether or not this identity is a consequence of an unreported breeding accident in the history of the MSM/MsJ colony.

Analysis of the expression of IGHJ genes showed little variation between strains, though interestingly, no strain shared the expression of the BALB/c IGHJ1*01 sequence. Analysis of IGHD gene expression was more informative. The NOD/ShiLtJ mice appear to carry an IGHD locus that is identical to that of the C57BL/6 strain. Not only does this analysis support the conclusions from the IGHV gene analysis, but it gives confidence in the inference process itself. The IGHD genes of the MSM strain were also similar to those of the C57BL/6 strain, with seven of the nine C57BL/6 IGHD sequences being shared. Against expectations, just five of the ten BALB/c IGHD sequences were shared by *M. m. domesticus*-derived LEWES mice, while 8 sequences were shared with the *M. m. musculus*-derived PWD strain. We cannot rule out the possibility that this analysis failed to identify SNVs in either the terminal 5’ or 3’ nucleotides of the IGHD sequences, though we believe this is unlikely. It has been shown that the inference process is often unable to accurately identify terminal nucleotides in AIRR-Seq data ^29^. On the other hand, the antibody repertoire of the mouse is strongly shaped by joining at sequences of short homology ^30^ and there may be strong evolutionary pressure maintaining, for example, the identity between the 3’ ends of the IGHD2 gene family and the 5’ ends of some IGHJ sequences, thereby preventing the development of variation in the terminal IGHD gene nucleotides.

The demonstration that NOD/ShiLtJ mice carry IGHV, IGHD and IGHJ loci that are so closely related to those of the C57BL/6 strain is intriguing and also quite unexpected, because no direct relationship between these strains has been reported. The NOD/ShiLtJ mouse was derived in the 1970s from cataract-prone CTS mice, which in turn were developed in the 1960s from outbred Swiss mice ^31, 32^. C57BL/6 mice, on the other hand, were developed in the 1920s by Clarence Little from the progeny of a pair of ‘fancy mice’ ^33, 34^. It is difficult to believe that the NOD and C57BL/6 loci could have arisen independently by the chance selection of unrelated outbred founder pairs. The nearly identical sets of IGHV genes in the C57BL/6 and NOD/ShiLtJ mice are more suggestive of introgression, with a C57BL/6-like locus being introduced into the ancestors of the modern NOD strain by outcrossing. This notion is further supported by the observation that NOD clusters together with C57BL/6, based on mitochondrial SNV, but is rather distant from the strain based on chromosome Y SNV ^28^. This hints of a contamination via the maternal line. Of note, this would not be the first reported breeding accident involving the NOD lineage ^35^.

If the IGHV and IGHD genes of the LEWES/EiJ, PWD/PhJ and CAST/EiJ mice are accepted as being broadly representative of the three major subspecies of the house mouse, then neither the BALB/c strain nor the C57BL/6 strain can be unequivocally linked with one or other of the three major subspecies of the house mouse. It may be that the IGH loci of these and other classical inbred strains have a mosaic structure, representing alternating blocks of genes that are potentially polymorphic within one of the three major mouse subspecies, or divergent between them. It is also possible that the IGH loci of these strains include haplotype blocks derived from other lineages of the house mouse, or even from other Mus species such as *M. spretus*. Other subspecies of *M. m. musculus* probably exist, and a number have been proposed, such as *M. m. bactrianus* and *M. m. gentilulus* ^36, 37^.

The divergence patterns of the IGH loci of the mouse strains reported here can be considered in the context of the allelic diversity that has been reported in humans. For example, a comparison of the two published fully sequenced human IGHV haplotypes revealed that, of the 68 non-redundant functional/ORF IGHV sequences identified across the two haplotypes, only 22 were observed in both, accounting for both allelic and structural variants ^18^. In addition, although the population genetics of the human IGH locus remains relatively uncertain ^18–20^, some indirect measures of diversity are available to us. One measure of diversity is provided by consideration of heterozygous loci in individuals who have been genotyped by the analysis of VDJ rearrangements. When heterozygosity was explored at 50 IGHV gene loci in 98 individuals, only 5 genes were heterozygous in more than 50% of individuals, and only 19 genes were heterozygous in more than 20% of individuals ^15^. However, it is notable that more extreme examples of IGHV heterozygosity have been reported in some populations; for example, a study of genomic DNA in 28 South Africans characterized >50 IGHV alleles in all but two individuals ^16^. This same study characterized 123 alleles that were not present in IMGT. Similarly, a study of 10 individuals from Papua New Guinea identified 17 previously unreported IGHV sequences; however, in each individual, all but two or three sequences had previously been reported from studies in Europe, America and Australia ^17^. Collectively, these data suggest that there are many alleles common across different populations, but in some cases, it is likely that some alleles will also be more or less frequent in, or even private to specific populations. Taken together, it seems likely that the level of IGH diversity in inbred laboratory mice could be more extensive than what has been observed in human studies conducted to date. This may be expected given that our data suggest that IG genes present in many of the strains in use today are likely to harbor variation originating from many different subspecies.

If we are to properly understand the antibody repertoires of the major laboratory strains, their IGH loci will all need to be separately investigated. Unfortunately, existing projects such as the Mouse Genome Project (MGP) are unlikely to provide the necessary data. The MGP is currently sequencing the genomes of 16 inbred mouse strains, but it is unlikely that such assemblies will lead to suitably reliable reference sequences for the IGH locus in the short term, particularly without targeted and focused efforts that attempt to deal with the complexity and associated technical issues of these regions. Thus, immediate advances in our knowledge of the IGH loci of inbred strains will likely come from the inference of genes from AIRR-seq data. The resulting lack of knowledge of non-coding regions, and the lack of positional data, may make it difficult to decipher relationships between some strains, but it should provide the basic information regarding germline genes that is needed for accurate repertoire studies. Indeed, from our reanalyses (Table 2) of data based on the novel germline sets reported here, it is clear that strain-specific data will be needed in many cases if we are to ensure precision in AIRR-seq studies of strains other than C57BL/6.

Deeper sequencing will also be required if a more complete documentation of the available repertoire of germline genes in any inbred strain is to be determined. Nevertheless, the partial documentation of the IGH loci reported here, and the strain differences seen are sufficient to raise the possibility that the IGHV locus may contribute to strain-related differences in mouse models of human disease. Allelic variants have been associated with differences in disease susceptibility of rats ^38^ and humans ^19^. IGHV sequence variability might also contribute to the differences that have been reported in the susceptibility of inbred mouse strains to both infectious ^39, 40^ and autoimmune diseases ^41^.

The discovery of striking differences in the number of apparently functional IGHV genes between the C57BL/6 and the BALB/c strains first raised the possibility that the continued use of a positional nomenclature system for IGHV genes could be problematic ^5^. The results presented here confirm that this is the case. The C57BL/6 locus is unable to serve as a map for IGHV genes from other strains, and this appears to be the case even for inbred strains that were shown in earlier SNV analysis to carry *M. m. musculus*-derived IGH loci. A non-positional nomenclature should therefore be developed. Attention should also be paid to the light chain loci carried by different mouse strains. We have shown by SNV analysis that the three major subspecies of the house mouse may all have contributed to the kappa loci of the major strains of inbred laboratory mice ^42^. If the kappa and lambda loci of laboratory mice are shown to have the same kind of strain to strain variability as we have shown here for the IGH locus, then a new non-positional nomenclature will be required for all the genes of the IG loci.

## METHODS

### Antibody gene repertoire sequencing

Whole dissected spleens, preserved in RNAlater, were obtained from female mice from Jackson Laboratories (https://www.jax.org) for five inbred strains (CAST/EiJ [JAX stock #000928], n=1; LEWES/EiJ [JAX stock #002798], n=1; PWD/PhJ [JAX stock #004660], n=1; MSM/MsJ [JAX stock #003719], n=1; NOD/ShiLtJ [JAX stock #001976], n=1). An individual mouse was studied from each strain, as the power of this kind of investigation comes from sequencing depth rather than from the investigation of biological replicates. Each VDJ rearrangement provides independent support for the presence of a gene segment in the genome.

Total RNA was extracted from a section of each spleen using the RNeasy Mini kit (Qiagen, Cat. No. 74104; Germantown, MD). For each strain, 5’ RACE first-strand cDNA synthesis was conducted using the SMARTer RACE cDNA Amplification Kit (Takara Bio, Cat. No. 634858; Mountain View, CA), with an input amount of 1 µg of RNA per sample. Rearranged VDJ-IgM amplicons were generated using a single IgM oligo positioned in the CH3 region of the mouse IGHM gene (5’-CAGATCCCTGTGAGTCACAGTACAC-3’; 10 µM), paired with a universal primer (Takara Bio; 5’-AAGCAGTGGTATCAACGCAGAGT-3’; 10 µM). First-strand cDNA for each strain was amplified using Thermo Fisher Phusion HF Buffer (Thermo Fisher, Cat. No. F530S; Waltham, MA) for 30 PCR cycles. Amplicons were run on 2% agarose gel for size confirmation. Final amplicons were used to generate SMRTbell template libraries and each library was sequenced across 2 SMRTcells on a Pacific Biosciences RSII system using P6/C4 chemistry and 240 minute movies (Pacific Biosciences; Menlo Park, CA).

### Data processing and germline gene inference

Reads from each RSII run were combined and processed using the CCS2 algorithm. CCS2 reads of Q30 or greater were processed with pRESTO ^43^ v0.5.1 as follows: 1) Reads without a universal primer were removed using a maximum primer match error rate of 0.2 and a maximum template-switch error rate of 0.5 to de-duplicate the reads; 2) Reads without an IgM primer were removed using a maximum primer match error rate of 0.2 and a maximum template-switch error rate of 0.5; 3) Duplicate sequences from the FASTQ files were removed using the default value of a maximum of 20 ambiguous nucleotides. Following pRESTO processing, IGHV gene assignments were noted after mapping reads to the IMGT germline database using IMGT/HighV-QUEST version 1.6.0 (20 June 2018) ^44^. Resulting IMGT summary output was processed using Change-O v0.4.3 ^45^. Reads representing incomplete or non-productive VDJ rearrangements were discarded. All sequences in each sample were assigned to clones using the distToNearest function in the SHazaM R package and the DefineClones function in Change-O ^45^. A single member of each clonal group was randomly selected for downstream analyses.

For germline IGHV gene inference for each strain, the sequences representing the strain’s clonal groups were first clustered based on the IGHV gene assignment and associated percent identity of the alignment to the nearest IMGT reference directory gene sequence. Sequences shorter than the 138 base pair (bp) length of the shortest reported functional IGHV sequence in IMGT were excluded from the analysis. Consensus IGHV gene sequences were then determined for each cluster using CD-HIT (cd-hit-est v4.6.8) ^46^ requiring that a given inference be represented by at least 0.1% of total clonal groups identified in the strain’s dataset. To be conservative, consensus sequences for each inferred germline were trimmed to the shortest sequence representative in the identified cluster. If the percent match identity to the closest IMGT germline gene sequence was <95%, then the alignments were manually inspected for evidence of chimeric PCR amplification ^47^.

An inferred germline IGHV sequence reference dataset was generated for each strain. Predicted germline IGHV gene databases from Q30 datasets were assessed using one iteration of IgDiscover v0.10 ^22^. For the analysis of each strain, inferred germline IGHV sequences generated from our clustering method were used as the IGHV gene database for IgDiscover. Changes made to the default configuration were as follows: 1) ‘race_g’ set to ‘true’ to account for a run of G nucleotides at the beginning of the sequence 2) ‘stranded’ set to ‘false’ because the forward primer was not always located at the 5’ end, and 3) ‘ignore_j’ set to ‘true’ to ignore whether or not a joining (J) gene had been assigned for a newly discovered IGHV gene.

All inferred germline sequences were compared, using BLAST (https://blast.ncbi.nlm.nih.gov/) ^48^, to sequences in the following databases: IMGT (http://www.imgt.org), VBASE2 (http://www.vbase2.org/), and the NCBI non-redundant nucleotide sequence collection. Germline sets were compared across strains using BLAT ^49^. Perfect matches between sequences of different strains required full length alignments of the query sequence at 100% identity; sequence length variation at the 3’ end of query and subject sequences was allowed. Calculations of mean IGHV sequence identities between strains were computed by taking the mean of the sequence identities for the best pair-wise hits of all genes in the smaller germline set of two strains being compared. For the comparison between IGHV sequences of NOD/ShiLtJ and MSM/MsJ, which had an equal number of inferred germline genes, we opted to present the mean sequence identity using the NOD/ShiLtJ germline set.

Germline IGHD genes were inferred for each strain by further analysis of the junction sequences from each strain’s clone group representatives. Junctional sequences as defined by IMGT/HighV-QUEST from the Change-O tables were compared against the full IMGT mouse IGHD reference directory (Release 201918-4; 9 May 2019). Reference directory genes that share identical coding sequences were grouped to a single entry for comparison to the VDJ junctions. For each reference IGHD gene all substrings of length 5 or more were compared to the all junction sub-sequences of the same length. Matches between the IGHD and junctional substrings were retained if the hamming distance was 2 or less, with the exception of rejecting the alignment if the pattern of mismatches fell into either of the following categories; the mismatches were in the first or final two positions of the substrings, or the third and fourth positions were both mismatched, or the third and fourth to last positions were both mismatched. These heuristics were implemented to avoid patterns of mismatch that arise from mis-alignment of non-template encoded N additions in circumstances where the IGHD has undergone exonuclease trimming. For each VDJ junction the set of IGHD substrings that maximised length and minimising mismatches were included in the analysis. Multiple IGHD substring matches were permitted per VDJ junction.

Counts for observed IGHD substring matches across each strain’s dataset of clone representations were tallied for analysis. Analysis considered all IGHD substring matches with fewer than 10 exonuclease removals and with minimum lengths of 10 nucleotides (or 9 for IGHD4-4). The presence of a reference IGHD was accepted if abundant, full-length matches were observed within a strain’s datasets (generally in excess of 10 observations). For reference gene-derived substrings that included mismatches, if the same mismatch was observed across 80% of the substrings for the IGHD gene then this was taken as putative evidence of the presence of SNVs in the germline gene. If the mismatches could be explained by mis-assignment of another IGHD which was already confirmed by full-length, perfect matches then the putative variant was dismissed. In the absence of such mis-assignment and with the support of the repeated detection of the SNV(s) associated with varied patterns of exonuclease removal, a new IGHD variant was defined. For strains with new IGHD variants, the substring analysis was repeated using the IMGT reference directory amended to replace the IMGT gene(s) with the novel variant(s). Final tabulations were made from these strain adjusted references.

IGHJ genes for each strain were inferred by analysis of the subset of clone representatives for each strain that had unmutated IGHV regions in order to reduce the likelihood of the IGHJ containing somatic point mutations. IGHJs that were more than 2 nucleotides shorter than the closest germline reference sequence were also excluded. For each strain’s dataset, IGHJ gene sequences were then clustered using CD-HIT, requiring exact matches. Consensus sequences from each cluster were then examined based on the number of supporting sequences. Within each strain, the top four clusters based on their observed frequency in the repertoire were taken to represent the IGHJ genes of that strain. Any additional clusters that had a frequency representing >10% of one of the top four IGHJ gene clusters were also further examined. In only one case did this occur (in CAST/EiJ); this sequence discarded because it was was determined to be a derivative of one of the top four clusters from this strain, and to include divergent bases originating from 5’ NP addition.

Finally, each strain’s dataset was re-aligned using strain-specific IGHV, IGHD and IGHJ references. Strain specific references included the sequences listed in Supplemental tables 2, 3 and 4. IMGT/HighV-QUEST was unable to be utilised for this analysis as it does not permit specification of the non-IMGT germline reference sets. Alignments were therefore performed with stand-alone IgBLAST ^50^ (version 1.14.0) with a reward score of +1 and a mismatch penalty of −3. Scoring was adjusted in this way due to length differences (283 - 303) for the inferred IGHVs due to conservative 3’ end trimming. The default +1/-1 scoring favours longer mismatched alignments, over shorter unmutated alignments, even when the shorter alignments are full length with respect to the inferred gene. The IgBLAST mouse auxiliary data file was amended to include the novel IGHJ2 variant from CAST/EiJ using the same offsets as IGHJ2*01. IgBLAST with the same parameters was also used to align that VDJ datasets against the IMGT reference directory to distinguish the contributions of the alignment algorithm and the germline reference.

**Additional References for Supplementary figures**

---

**Supplementary figure 1.** Single nucleotide variant (SNV) data from the IGHV gene region (chr12:114700000-117270000) suggest common subspecies origins for IGHV genomic haplotypes of inbred and wild-derived laboratory mouse strains. This figure depicts the predicted subspecies origins (*Mus musculus domesticus*; *M. m. musculus*; *M. m. castaneous*) of IGHV haplotypes in the six strains analyzed in the present study. Here, we consider the relationships between inferred germline IGHV gene sets of these strains in the context of these predicted subspecific origins. Data presented in this figure were obtained from the Mouse Phylogeny Viewer ^1^ (https://msub.csbio.unc.edu/, based on previously published whole-genome SNV data ^2^. The bars above each haplotype depict the locations of “diagnostic” SNVs used to make subspecies determinations.

**Supplementary figure 2.** Counts of identified clones representing inferred germline sequences from each strain sequenced in this study. Additional information, including the nucleotide sequences of each inferred germline presented in these plots, can be found in Supplemental Table 2.

**Supplementary figure 3.** Boxplots depicting the numbers of clones representing each inferred IGHV germline sequence in CAST/EiJ, LEWES/EiJ, MSM/MsJ, and PWD/PhJ, respectively. Data for each strain are partitioned by whether the inferred sequence was supported by IgDiscover. With only a few exceptions, the inferred IGHV sequences in each strain lacking support by IgDiscover (“No”) were represented by fewer clones in our analysis on average than those that were supported by IgDiscover (“Yes”).

**Supplementary figure 4.** Violin plots depicting the best percent match identities among inferred IGHV germline sets for each pair-wise strain comparison. Individual points within each violin plot represent the best percent identity value of an IGHV gene segment as compared to all genes in the other strain to which it was compared.

**Supplementary table 1.** Per-strain AIRR-seq library summary statistics for CAST/EiJ, LEWES/EiJ, MSM/MsJ, PWD/PhJ, and NOD/ShiLtJ.

**Supplementary table 2.** Complete database of inferred germline IGHV sequences from CAST/EiJ, LEWES/EiJ, MSM/MsJ, PWD/PhJ, and NOD/ShiLtJ.

**Supplementary table 3.** Complete dataset of inferred germline IGHD sequences from CAST/EiJ, LEWES/EiJ, MSM/MsJ, PWD/PhJ, and NOD/ShiLtJ.

**Supplementary table 4.** Complete dataset of inferred germline IGHJ sequences from CAST/EiJ, LEWES/EiJ, MSM/MsJ, PWD/PhJ, and NOD/ShiLtJ.

## REFERENCES

1. Morse HC. Origins of inbred mice. Academic Press: New York, 1978.

2. Alt FW, Baltimore D. Joining of immunoglobulin heavy chain gene segments: implications from a chromosome with evidence of three D-JH fusions. Proc Natl Acad Sci USA 1982; 79: 4118–4122.

3. Leder P, Honjo T, Packman S, Swan D, Nau M, Norman B. The organization and diversity of immunoglobulin genes. Proc Natl Acad Sci USA 1974; 71: 5109–5115.

4. Potter M. Antigen-binding myeloma proteins of mice. Adv Immunol 1977; 25: 141–211.

5. Collins AM, Wang Y, Roskin KM, Marquis CP, Jackson KJ. The mouse antibody heavy chain repertoire is germline-focused and highly variable between inbred strains. Philos Trans R Soc Lond B Biol Sci 2015; 370.

6. Johnston CM, Wood AL, Bolland DJ, Corcoran AE. Complete sequence assembly and characterization of the C57BL/6 mouse Ig heavy chain V region. J Immunol 2006; 176: 4221–4234.

7. Riblet R. Immunoglobulin heavy chain genes in the mouse. In: Honjo T, Alt FW, Neuberger M (eds). Molecular biology of B cells. London: Elsevier Academic Press; 2004, pp 19–26.

8. Retter I, Chevillard C, Scharfe M, et al. Sequence and characterization of the Ig heavy chain constant and partial variable region of the mouse strain 129S1. J Immunol 2007; 179: 2419–2427.

9. Retter I, Althaus HH, Munch R, Muller W. VBASE2, an integrative V gene database. Nucleic Acids Res 2005; 33: D671–674.

10. Giudicelli V, Duroux P, Ginestoux C, et al. IMGT/LIGM-DB, the IMGT comprehensive database of immunoglobulin and T cell receptor nucleotide sequences. Nucleic Acids Res 2006; 34: D781–784.

11. Matthyssens G, Rabbitts TH. Structure and multiplicity of genes for the human immunoglobulin heavy chain variable region. Proc Natl Acad Sci U S A 1980; 77: 6561–6565.

12. Matsuda F, Ishii K, Bourvagnet P, et al. The complete nucleotide sequence of the human immunoglobulin heavy chain variable region locus. J Exp Med 1998; 188: 2151–2162.

13. Boyd SD, Gaeta BA, Jackson KJ, et al. Individual variation in the germline Ig gene repertoire inferred from variable region gene rearrangements. J Immunol 184: 6986–6992.

14. Glanville J, Zhai W, Berka J, et al. Precise determination of the diversity of a combinatorial antibody library gives insight into the human immunoglobulin repertoire. Proc Natl Acad Sci USA 2009; 106: 20216–20221.

15. Gidoni M, Snir O, Peres A, et al. Mosaic deletion patterns of the human antibody heavy chain gene locus shown by Bayesian haplotyping. Nature communications 2019; 10: 628.

16. Scheepers C, Shrestha RK, Lambson BE, et al. Ability to develop broadly neutralizing HIV-1 antibodies is not restricted by the germline Ig gene repertoire. J Immunol 2015; 194: 4371–4378.

17. Wang Y, Jackson KJ, Gaeta B, et al. Genomic screening by 454 pyrosequencing identifies a new human IGHV gene and sixteen other new IGHV allelic variants. Immunogenetics 2011; 63: 259–265.

18. Watson CT, Steinberg KM, Huddleston J, et al. Complete haplotype sequence of the human immunoglobulin heavy-chain variable, diversity, and joining genes and characterization of allelic and copy-number variation. Am J Hum Genet 2013; 92: 530–546.

19. Avnir Y, Watson CT, Glanville J, et al. IGHV1-69 polymorphism modulates anti-influenza antibody repertoires, correlates with IGHV utilization shifts and varies by ethnicity. Sci Rep 2016; 6: 20842.

20. Watson CT, Glanville J, Marasco WA. The Individual and population genetics of antibody immunity. Trends Immunol 2017; 38: 459–470.

21. Ohlin M, Scheepers C, Corcoran M, et al. Inferred allelic variants of immunoglobulin receptor genes: a system for their evaluation, documentation, and naming. Front Immunol 2019; 10: 435.

22. Corcoran MM, Phad GE, Vazquez Bernat N, et al. Production of individualized V gene databases reveals high levels of immunoglobulin genetic diversity. Nature communications 2016; 7: 13642.

23. Gadala-Maria D, Gidoni M, Marquez S, et al. Identification of subject-specific immunoglobulin alleles from expressed repertoire sequencing data. Front Immunol 2019; 10: 129.

24. Ralph DK, Matsen IV FA. Per sample immunoglobulin germline inference from B cell receptor deep sequencing data. arXiv:171105843v2 [q-bioPE] 2018.

25. Zhang W, Wang IM, Wang C, et al. IMPre: An accurate and efficient software for prediction of T- and B-cell receptor germline genes and alleles from rearranged repertoire data. Front Immunol 2016; 7: 457.

26. Jouvin-Marche E, Morgado MG, Leguern C, Voegtle D, Bonhomme F, Cazenave PA. The mouse Igh-1a and Igh-1b H chain constant regions are derived from two distinct isotypic genes. Immunogenetics 1989; 29: 92–97.

27. Collins AM, Jackson KJL. On being the right size: antibody repertoire formation in the mouse and human. Immunogenetics 2018; 70: 143–158.

28. Yang H, Wang JR, Didion JP, et al. Subspecific origin and haplotype diversity in the laboratory mouse. Nat Genet 2011; 43: 648–655.

29. Thornqvist L, Ohlin M. Critical steps for computational inference of the 3’-end of novel alleles of immunoglobulin heavy chain variable genes - illustrated by an allele of IGHV3-7. Mol Immunol 2018; 103: 1–6.

30. Feeney AJ. Predominance of VH-D-JH junctions occurring at sites of short sequence homology results in limited junctional diversity in neonatal antibodies. J Immunol 1992; 149: 222–229.

31. Makino S, Kunimoto K, Muraoka Y, Mizushima Y, Katagiri K, Tochino Y. Breeding of a non-obese, diabetic strain of mice. Jikken Dobutsu 1980; 29: 1–13.

32. Mullen Y. Development of the Nonobese diabetic mouse and contribution of animal models for understanding type 1 diabetes. Pancreas 2017; 46: 455–466.

33. Beck JA, Lloyd S, Hafezparast M, et al. Genealogies of mouse inbred strains. Nat Genet 2000; 24: 23–25.

34. Staats J. The laboratory mouse. In: Green EL (ed). Biology of the laboratory mouse. New York: McGraw-Hill; 1966, pp 1–9.

35. Prochazka M, Serreze DV, Frankel WN, Leiter EH. NOR/Lt mice: MHC-matched diabetes-resistant control strain for NOD mice. Diabetes 1992; 41: 98–106.

36. Suzuki H, Nunome M, Kinoshita G, et al. Evolutionary and dispersal history of Eurasian house mice Mus musculus clarified by more extensive geographic sampling of mitochondrial DNA. Heredity (Edinb) 2013; 111: 375–390.

37. Suzuki H, Yakimenko LV, Usuda D, Frisman LV. Tracing the eastward dispersal of the house mouse, Mus musculus. Genes and environment 2015; 37: 20.

38. Dhande IS, Cranford SM, Zhu Y, et al. Susceptibility to hypertensive renal disease in the Spontaneously Hypertensive rat is influenced by 2 loci affecting blood pressure and immunoglobulin repertoire. Hypertension 2018; 71: 700–708.

39. Caron J, Loredo-Osti JC, Laroche L, Skamene E, Morgan K, Malo D. Identification of genetic loci controlling bacterial clearance in experimental Salmonella enteritidis infection: an unexpected role of Nramp1 (Slc11a1) in the persistence of infection in mice. Genes Immun 2002; 3: 196–204.

40. Swihart K, Fruth U, Messmer N, et al. Mice from a genetically resistant background lacking the interferon gamma receptor are susceptible to infection with Leishmania major but mount a polarized T helper cell 1-type CD4+ T cell response. J Exp Med 1995; 181: 961–971.

41. Hannestad K, Scott H. The MHC haplotype H2b converts two pure nonlupus mouse strains to producers of antinuclear antibodies. J Immunol 2009; 183: 3542–3550.

42. Collins AM, Watson CT. Immunoglobulin light chain gene rearrangements, receptor editing and the development of a self-tolerant antibody repertoire. Front Immunol 2018; 9: 2249.

43. Vander Heiden JA, Yaari G, Uduman M, et al. pRESTO: a toolkit for processing high-throughput sequencing raw reads of lymphocyte receptor repertoires. Bioinformatics 2014; 30: 1930–1932.

44. Alamyar E, Giudicelli V, Duroux P, Lefranc MP. Antibody V and C domain sequence, structure, and interaction analysis with special reference to IMGT. Methods Mol Biol 2014; 1131: 337–381.

45. Gupta NT, Vander Heiden JA, Uduman M, Gadala-Maria D, Yaari G, Kleinstein SH. Change-O: a toolkit for analyzing large-scale B cell immunoglobulin repertoire sequencing data. Bioinformatics 2015; 31: 3356–3358.

46. Huang Y, Niu B, Gao Y, Fu L, Li W. CD-HIT Suite: a web server for clustering and comparing biological sequences. Bioinformatics 2010; 26: 680–682.

47. Meyerhans A, Vartanian JP, Wain-Hobson S. DNA recombination during PCR. Nucleic Acids Res 1990; 18: 1687–1691.

48. Altschul SF, Gish W, Miller W, Myers EW, Lipman DJ. Basic local alignment search tool. J Mol Biol 1990; 215: 403–410.

49. Kent WJ. BLAT--the BLAST-like alignment tool. Genome Res 2002; 12: 656–664.

50. Ye J, Ma N, Madden TL, Ostell JM. IgBLAST: an immunoglobulin variable domain sequence analysis tool. Nucleic Acids Res 2013.

## REFERENCES

1. Wang JR, de Villena FP, McMillan L. Comparative analysis and visualization of multiple collinear genomes. BMC Bioinformatics 2012; 13: S13.

2. Yang H, Wang JR, Didion JP et al. Subspecific origin and haplotype diversity in the laboratory mouse. Nat Genet 2011; 43: 648–655.

