## Supplementary figures and images for "A comparison of immunoglobulin IGHV, IGHD and IGHJ genes in wild-derived and classical inbred mouse strains"

### Supplmental figure 1

#### IGHV region (chr12:114700000-117270000)

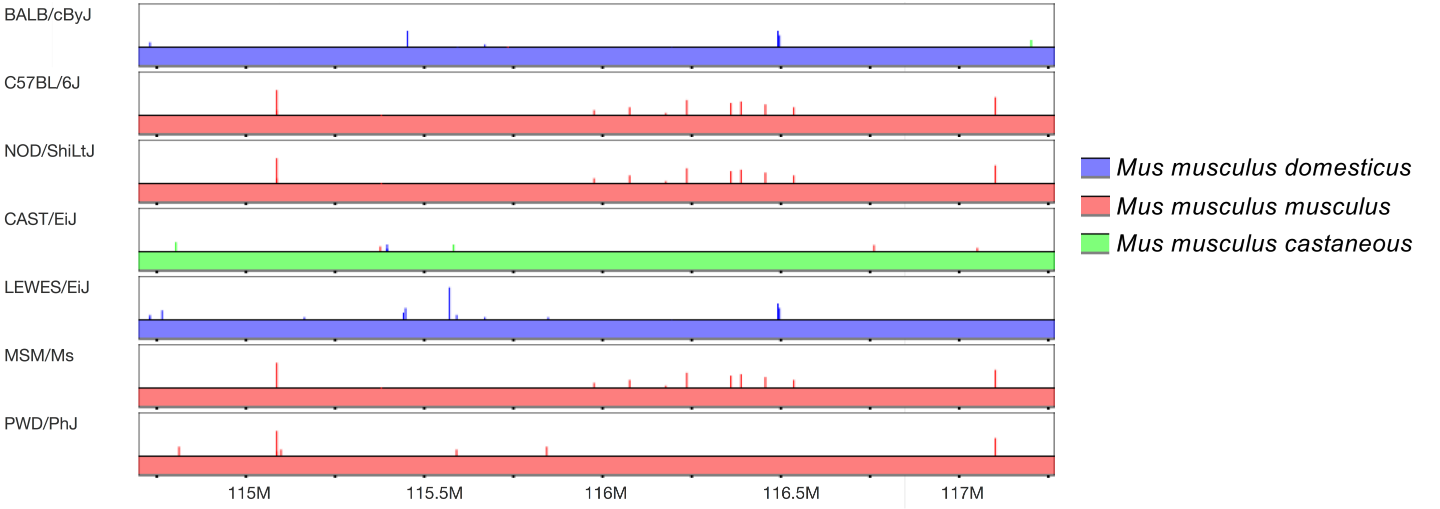

### Supplmental figure 2

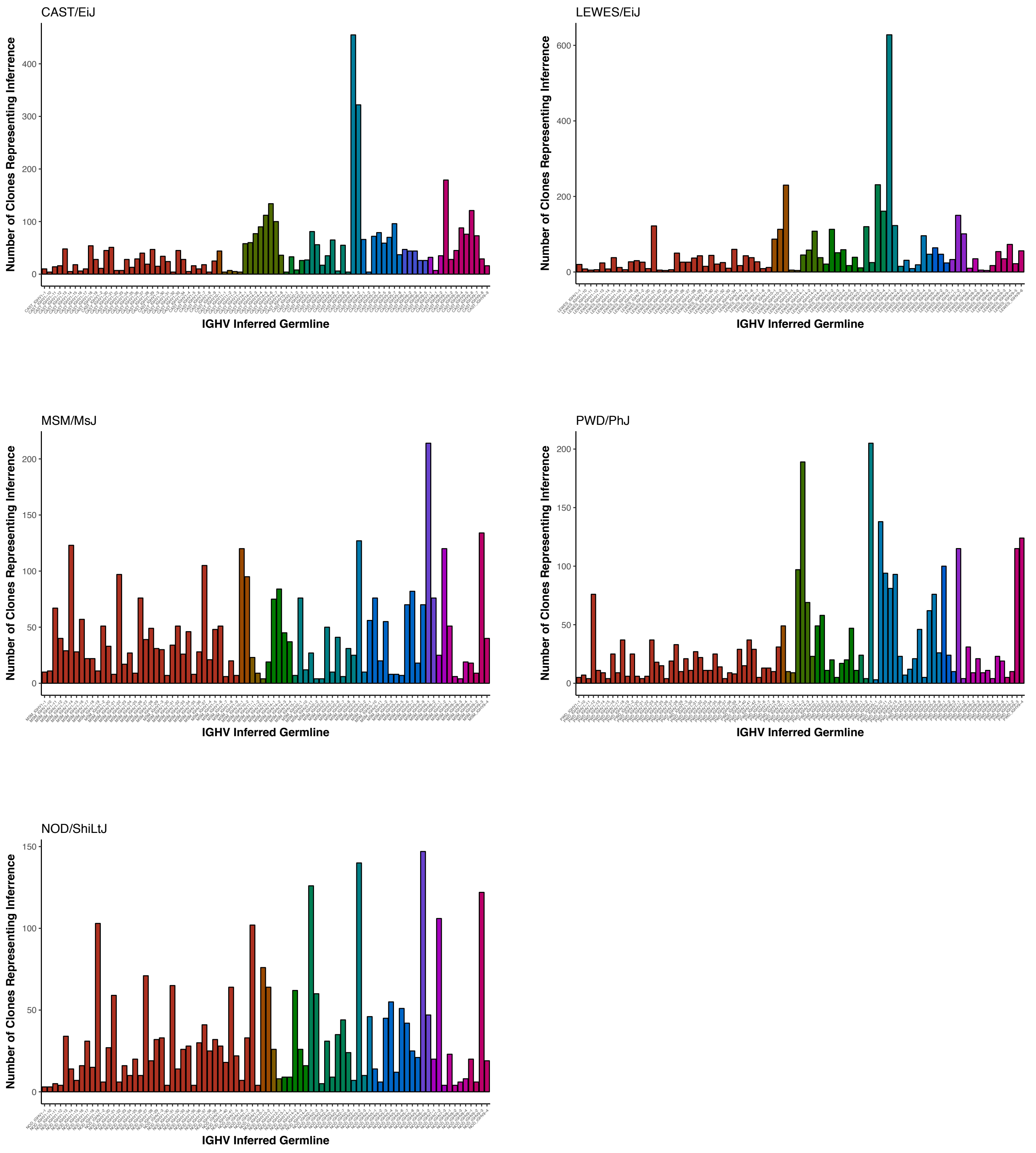

### Supplmental figure 3

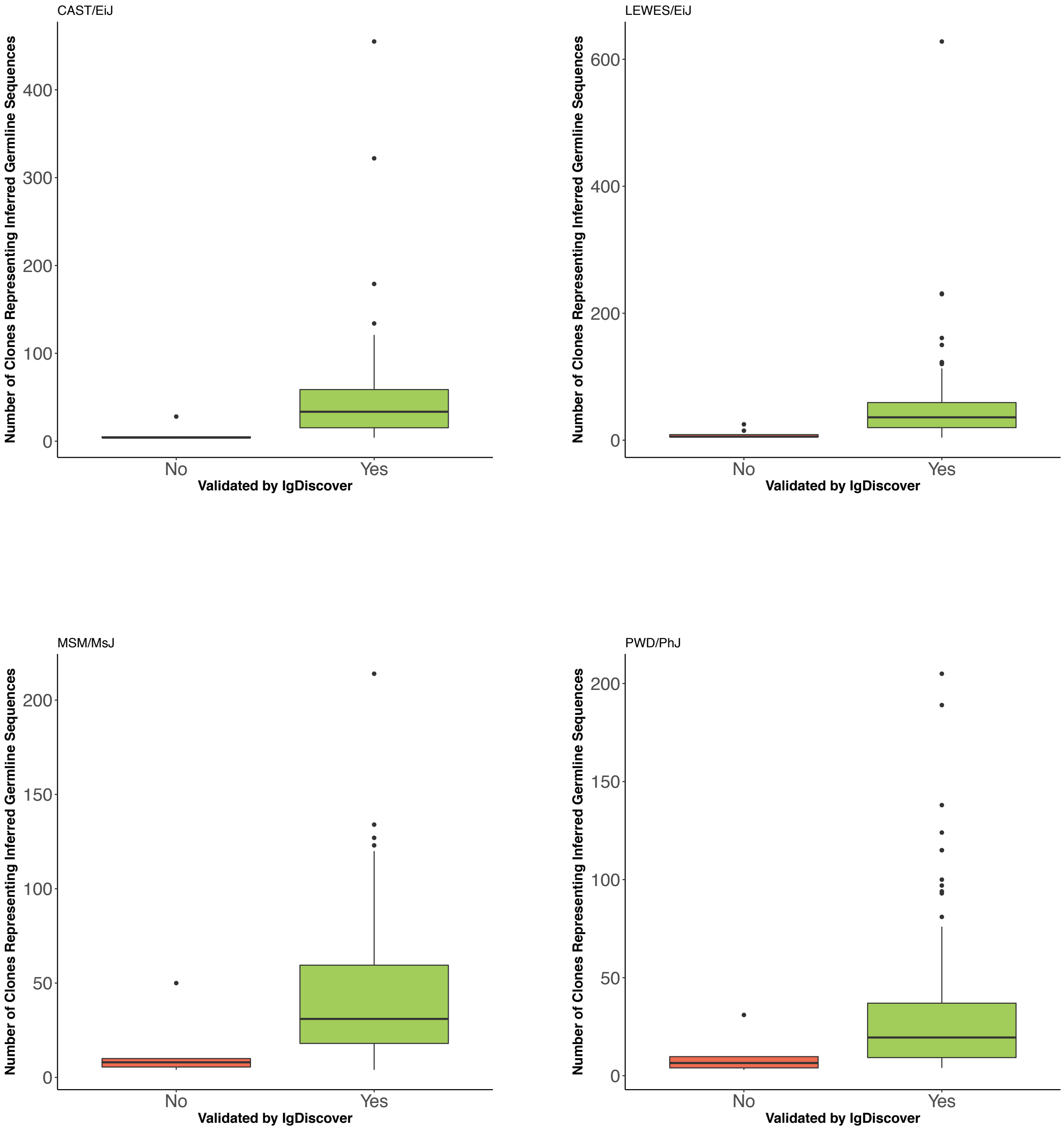

### Supplmental figure 4

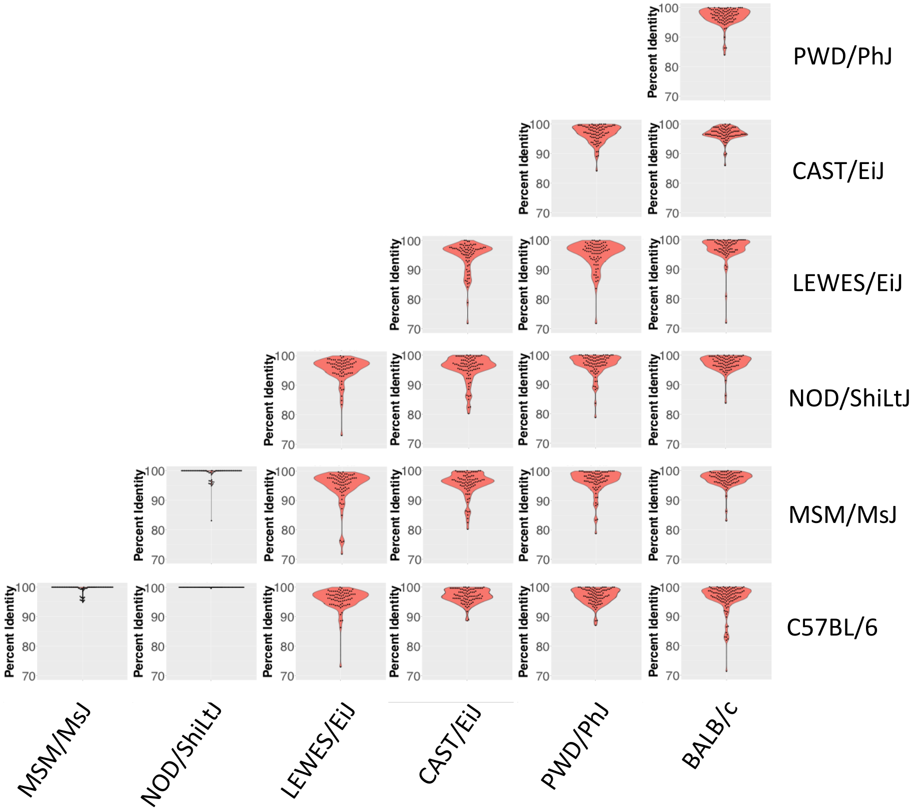
